# No evidence for phylostratigraphic bias impacting inferences on patterns of gene emergence and evolution

**DOI:** 10.1101/060756

**Authors:** Tomislav Domazet-Lošo, Anne-Ruxandra Carvunis, M.Mar Albà, Martin Sebastijan Šestak, Robert Bakarić, Rafik Neme, Diethard Tautz

## Abstract

Phylostratigraphy is a computational framework for dating the emergence of sequences (usually genes) in a phylogeny. It has been extensively applied to make inferences on patterns of genome evolution, including patterns of disease gene evolution, ontogeny and *de novo* gene origination. Phylostratigraphy typically relies on BLAST searches along a species tree, but new simulation studies have raised concerns about the ability of BLAST to detect remote homologues and its impact on phylostratigraphic inferences. These simulations called into question some of our previously published work on patterns of gene emergence and evolution inferred from phylostratigraphy. Here, we re-assessed these simulations and found major problems including unrealistic parameter choices, irreproducibility, statistical flaws and partial representation of results. We found that, even with a possible overall BLAST false negative rate between 5-15%, the large majority (≥74%) of sequences assigned to a recent evolutionary origin by phylostratigraphy is unaffected by technical concerns about BLAST. Where the results of the simulations did cast doubt on our previous findings, we repeated our analyses but now excluded all questionable sequences. The originally described patterns remained essentially unchanged. These new analyses strongly support our published inferences, including: genes that emerged after the origin of eukaryotes are more likely to be expressed in the ectoderm than in the endoderm or mesoderm in *Drosophila*, and the *de novo* emergence of protein-coding genes from non-genic sequences occurs through proto-gene intermediates in yeast. We conclude that BLAST is an appropriate and sufficiently sensitive tool in phylostratigraphic analysis.

## Introduction

Correlating the emergence of particular sequences with molecular and phenotypic features is one way to harness the information that we obtain from genome sequencing projects. Phylostratigraphy is a framework in which this can be done in a phylogeny aware context (Domazet-Lošo et al. 2007). Starting from the genome of a focal species, phylostratigraphy infers the emergence of novel sequences at a particular phylogenetic node, usually by using the similarity search algorithm BLAST (Altschul et al. 1990) on a set of genomes that represent the nodes. Each sequence in the focal genome is thereby assigned an “evolutionary age” corresponding to the most distant node in the phylogeny where BLAST could detect a homologue for this sequence. This age classification, also referred to as “phylostrata” or “conservation level” classification, enables to distinguish younger sequences, for which homologues can only be found in closely related species (often called orphans or taxonomically restricted), from older sequences that are conserved in very distant species (Tautz and Domazet-Lošo 2011). While phylostratigraphy is a general evolutionary framework that in theory applies to any type of sequence, it has mostly been exploited to study the evolution of novel genes and open reading frames (ORFs).

It is important to note that novel genes can evolve through two different mechanisms. One is *de novo* evolution, which has only relatively recently been recognized as an important mechanism for evolution of novel genes (Levine et al. 2006; Zhou et al. 2008; Heinen et al. 2009; Knowles and McLysaght 2009; Toll-Riera et al. 2009; Carvunis et al. 2012; Neme and Tautz 2014). The other is rapid divergence from existing genes, for instance due to adaptation to a new function (Domazet-Lošo and Tautz 2003). The concept of phylostratigraphy was originally based on this latter mechanism and proposed the idea of a punctuated evolution of protein-coding genes and their descendant families (Domazet-Lošo et al. 2007; Domazet-Lošo and Tautz 2010a). Punctuated evolution assumes that a gene originates by duplication from an existing gene followed by fast divergence, likely due to a new adaptation, with a subsequent slow-down in sequence evolution. Such slow evolving orphan genes were first detected in *Drosophila,* and they were proposed to represent lineage-specific adaptations (Domazet-Lošo and Tautz 2003). Hence, the shifts in sequence space generated by phases of fast evolution after gene duplication are indicators of a new adaptive function and phylostratigraphy aims to trace such events and to statistically correlate them to biological patterns (Domazet-Lošo and Tautz 2010b; Quint et al. 2012; Mendoza et al. 2013; Šestak et al. 2013; Šestak and Domazet-Lošo 2015; Drost et al. 2016).

*De novo* emergence from a previously non-genic sequence can be equally detected by phylostratigraphy. For a long time, *de novo* emergence was considered to be very unlikely (Tautz 2014) and had therefore initially not been seriously considered as a model of origin of orphan genes (Domazet-Lošo and Tautz 2003). However, it is now clear that *de novo* gene birth is in fact another important process that can be traced by phylostratigraphy, in particular among closely related species (Tautz and Domazet-Lošo 2011). Accordingly, phylostratigraphy has also been used in later studies specifically focusing on the patterns and mechanisms of *de novo* evolution (Carvunis et al. 2012; Abrusán 2013; Neme and Tautz 2013).

Although BLAST is very powerful in detecting homologues in large databases, it has known limitations when sequences are highly diverged. In particular, BLAST has problems to detect remote homologues of short and fast-evolving sequences (Elhaik et al. 2006; Moyers and Zhang 2015). These limitations do not much affect evolutionary inferences related to punctuated evolution of proteins and their descendant families, where the existence of possible remote homologues is not the primary question (Domazet-Lošo et al. 2007; Domazet-Lošo and Tautz 2010a). If anything, BLAST could be too sensitive in this context, and find an older origin for a protein, although it has gone through a recent shift in sequence space. For example, transcription factors that have arisen to regulate a specific function in a young lineage may become placed into a much older node because of a match within their DNA binding domain (Capra et al. 2013). BLAST could also overestimate a protein’s evolutionary age by yielding spurious hits that do not reflect true homology. On the other hand, the difficulty of BLAST searches to find remote homologues could be problematic in the context of making cases for true *de novo* gene emergence, versus fast divergence from an ancient gene (Schlötterer 2015). Ancient genes that have diverged too much for BLAST to detect them in the genomes of distant species may then be erroneously categorized as too young by phylostratigraphy. These BLAST limitations have motivated the development of further refined search methods, such as PSI-BLAST (Altschul et al. 1997), HHMER3 (Finn et al. 2011) or HHblits (Remmert et al. 2012). Although these refined methods can detect more remote homologues, they are computationally more costly, require similarity profiles from well-populated gene families and are therefore less generally applicable. Hence, BLAST remains the workhorse for obtaining initial phylostratigraphic information and it is therefore important to understand its advantages, as well as its limitations and possible error margins.

In an attempt to estimate the false negative error rate of BLAST and its impact on evolutionary inferences, Elhaik et al. (2006) simulated DNA sequence evolution and used BLAST to look for homologues of these simulated sequences. They found in these simulations that fast-evolving DNA sequences tended to appear younger than they were, and suggested that the “Inverse Relationship Between Evolutionary Rate and Age of Mammalian Genes’ ‘ previously reported (Albà and Castresana 2005) may have been an artifact. This suggestion was rapidly refuted when Albà and Castresana (2007) pointed out a problem in the simulation framework used by Elhaik et al. (2006). BLAST uses a two-step search algorithm that starts by finding matches on short motifs and extending the alignment based on these (Altschul et al. 1990). Proteins that evolve homogeneously along their whole sequence are thus more difficult to trace than proteins that include at least one or more slowly evolving domains. Real proteins fall mostly into this latter class, allowing BLAST to find homologues even when the rest of a protein sequence evolves very fast. Therefore, Albà and Castresana (2007) argued that simulating protein evolution to assess the power of BLAST needs to take natural among-site rate heterogeneity into account.

Using this controlled approach, Albà and Castresana (2007) have shown that less than 5% of simulated homologues of mammalian genes are misclassified as recently evolved (i.e. too young) when rate heterogeneity is taken into account. The discussion on the potential biases introduced by false negatives in BLAST searches was therefore considered resolved (Tautz and Domazet-Lošo 2011). Using an orthogonal approach, Carvunis et al. (2012) have estimated that only 5% of ORFs appearing young in phylostratigraphy were likely to be misclassified due to false negatives in BLAST searches. This estimate was obtained by searching the entire non-redundant protein sequence database of NCBI for potential homologues of ORFs classified young in a phylostratigraphy of Ascomycota fungi.

Still, the question re-emerged recently when Moyers and Zhang (2015); Moyers and Zhang (2016) sought to quantify the power of BLAST to detect remote homologues once again, and to assess the possible implications for trends and patterns inferred from phylostratigraphic analysis. The first study (Moyers and Zhang 2015) criticizes Albà and Castresana (2007)’s work by stating that the rate heterogeneity models used in this study were derived from only 14 genes and may not have been typical. Hence, they used a much larger set of genes derived from *Drosophila melanogaster* and calculated among-site rate heterogeneity and average divergence rate for each gene based on an alignment among 12 *Drosophila* species. These actual genes and their associated divergence rates were then used to simulate their possible ancestors at the origin of life and ask which percentage of such ancestors can be traced by BLAST. They find that BLAST makes an incorrect assignment for 14% of the sequences simulated. According to their follow up analyses, these potential 14% of errors may have impacted two previously published evolutionary inferences: a peak of new gene origination in the common ancestor of bilateria, and a non-random age distribution of genes expressed in ectoderm, mesoderm and endoderm during *Drosophila* development (Domazet-Lošo et al. 2007). Moyers and Zhang (2015) also claimed that false negative errors of BLAST may explain another previously published inference according to which human disease genes tend to be ancient (Domazet-Lošo and Tautz 2008), although they do not provide an estimated error rate for human-centered phylostratigraphy.

In their second paper, Moyers and Zhang (2016) addressed the question of *de novo* evolution of genes in yeast species. They choose as title for this paper "Evaluating phylostratigraphic evidence for widespread *de novo* gene birth in genome evolution", but they address mostly a different question, namely the significance of observed trends in recently evolved yeast ORFs. Starting from protein sequence alignments between yeast species closely related to the focal species *Saccharomyces cerevisiae*, Moyers and Zhang (2016) measured among-site rate heterogeneity and average divergence rates, and simulated possible ancestors throughout the phylogeny based on the measured rates. They report that BLAST missed 11% of the simulated ancient homologues, i.e. more than in the original study (Carvunis et al. 2012) had reported. They show that the corresponding ORFs, which may erroneously appear young in phylostratography, despite potentially being ancient, share many physical and functional properties with the ORFs deemed young by Carvunis et al. (2012) and Abrusán et al. (2013). Based on these observations, Moyers and Zhang (2016) question the validity of genome-wide phylostratigraphic analyses for deriving models of *de novo* gene birth.

In summary, Moyers and Zhang (2015); Moyers and Zhang (2016) have revived several important technical and conceptual issues pertaining to an older debate on the limitations of BLAST (Albà and Castresana 2005; Elhaik et al. 2006; Albà and Castresana 2007). Here, we argue that Moyers and Zhang’s (2015); Moyers and Zhang’s 2016) simulations have underestimated the power of BLAST in phylostratigraphy. We show that the previously published inferences on gene emergence and evolution that were questioned by Moyers and Zhang (2015); Moyers and Zhang (2016) are in fact robust to BLAST limitations, even if error rates were as estimated by Moyers and Zhang (2015); Moyers and Zhang (2016). Finally, we clarify several points that were misinterpreted or misrepresented by Moyers and Zhang. We conclude that the alleged evidence for a systematic phylostratigraphic bias cannot be reproduced.

## Results

### The power of BLAST in phylostratigraphic analysis

The simulations performed by Moyers and Zhang suggested that up to 14% of *Drosophila melanogaster* sequences (2015) and up to 11% of *Saccharomyces cerevisiae* sequences (2016) may erroneously appear to have originated recently due to the limitations of BLAST. While this is a higher fraction than the one found by Albà and Castresana (2007), it is no reason to claim an "extreme" problem of age underestimation. Regardless, we argue that Moyer and Zhang’s estimates are likely to be exaggerated due to several technical issues.

First, the simulations use the actual gene sets and their corresponding divergence rates obtained from *Drosophila* and yeast alignments to evolve them in the simulations, i.e. they retain features that are inherent to these gene sets, rather than starting from simulated model sequences. Second, they do not use the same substitution matrix as the subsequent BLAST analysis, which could influence the outcome in untested ways. Third, dipteran insects, including *Drosophila*, are known to evolve about three times faster than other insects or vertebrates (Savard et al. 2006), which might have inflated the BLAST detection error in Moyers and Zhang (2015). Finally, the phylostratigraphy methodology used by Moyers and Zhang (2016) to search for remote homologues among their simulated yeast sequences is less sensitive than the one deployed in the original analysis of real sequences (Carvunis et al. 2012). In the original analyses, the authors assigned to each ORF sequence the conservation level of its most conserved paralogue, in an effort to avoid underestimating conservation (Carvunis et al. 2012). Moyers and Zhang (2016) did not implement this “oldest paralogue age” approach except in a single analysis, for which they did not report the corresponding BLAST false negative rate. Furthermore, where Moyers and Zhang (2016) used the program BLASTP, Carvunis et al. (2012) used three BLAST programs: BLASTP, TBLASTX, TBLASTN. The use of three BLAST programs necessarily results in a lower false negative rate than the use of a single program. This was noted by Moyers and Zhang but they chose not to take it into account, because "realistic simulation of codon sequence evolution is difficult" (quote from (Moyers and Zhang 2016)). It is thus evident that the false negative rates of the original phylostratigraphic analyses must be lower than those estimated by Moyers and Zhang.

We also note that Moyers and Zhang (2016) misinterpreted Abrusán (2013) by stating he ‘used Carvunis et al.’s data to examine a number of additional gene properties that he proposed to reflect the gradual genetic integrations of *de novo* genes into cellular networks or maturation of protein structures’ (quote from (Moyers and Zhang 2016)). However, Abrusán (2013) only used the classification of very young ORFs from Carvunis et al. (‘proto-genes’) but drew from the orthology classification provided by Wapinski et al. (2007) to classify all more conserved genes, which constitute the majority of annotated ORFs in the *S. cerevisiae* genome. The false negative rate associated with the methodology used by Abrusán (2013) was not estimated by Moyers and Zhang (2016).

Moyers and Zhang (2015) claimed that their estimates of BLAST detection errors are conservative, in particular due to not taking into account variations in rate heterogeneities across time. Such changes are indeed well known in phylogenetic analysis under the term covarion pattern of protein evolution (Penny et al. 2001). Moyers and Zhang (2015) simulate such a covarion pattern to assess BLAST performance in an attempt to provide an even more realistic framework of protein evolution. They find that BLAST performs indeed less well under these conditions, with up to 67% error rate in finding the oldest assignments. However, to obtain such a high rate of misplacement, they had to assume unrealistic parameters. This should already be evident from the fact that such a high misplacement rate is not compatible with real data, since most genes are actually mapped to the basal nodes in all phylostratigraphies (e.g. Domazet-Lošo and Tautz 2008; Tautz and Domazet-Lošo 2011). In their covarion model they shuffle over time the rates of up to 5% of sites per 50My and state that shuffling 1% of sites per 50My is a "tiny amount of covarion evolution" (quote from (Moyers and Zhang 2015)). However, when 2,500My of evolution are simulated, 1% per 50My amounts to 50% of the protein in total. Actual covarion proportions in old proteins were found to be around 10% (Wang et al. 2009). Hence, even 1% of sites per 50My is already beyond the realistic parameter space, let alone the 5% where they find the highest error rate. Even at the exaggerated 1% rate, the BLAST error is only around 18% (Table 2 in (Moyers and Zhang 2015)). The actual interpretation should therefore be that BLAST, when used in the phylostratigraphic framework, is very robust with respect to the rate heterogeneities found in real data.

Independent of whether the performance of BLAST to detect remote homologues results in 5% (Albà and Castresana 2007), 11% (Moyers and Zhang 2016) or 14% (Moyers and Zhang 2015) false negatives, it is clear that the large majority of assignments in phylostratigraphic studies is still not in doubt. Using real data, phylostratigraphy analyses have revealed that comparatively large numbers of genes lack clear remote homologues and many can only be assigned to the most recent nodes (Moyers and Zhang 2015; Moyers and Zhang 2016). Here, we updated the *Drosophila* phylostratigraphy for over 13K *Drosophila melanogaster* real sequences (supplementary table S1). We then compared the simulated and the real data obtained for 6,629 of these sequences, where simulated age assignments were available (supplementary table S1). The resulting distributions show that the large number of sequences lacking remote homologues in real data cannot be recapitulated by Moyers and Zhang’s simulations (Figure 1A). Similarly in yeast, ~40% of 5,878 *S. cerevisiae* ORFs for which both real (Carvunis et al. 2012) and simulated (Moyers and Zhang 2016) age data are available lack homologues in the distant species *S. pombe*, in stark contrast with the 11% estimate of misplaced ORFs by Moyers and Zhang (2016). The distributions of simulated versus actual data are again qualitatively very different (Figure 1B). Hence, the patterns found for real data are robust to BLAST errors.

**Figure 1:**
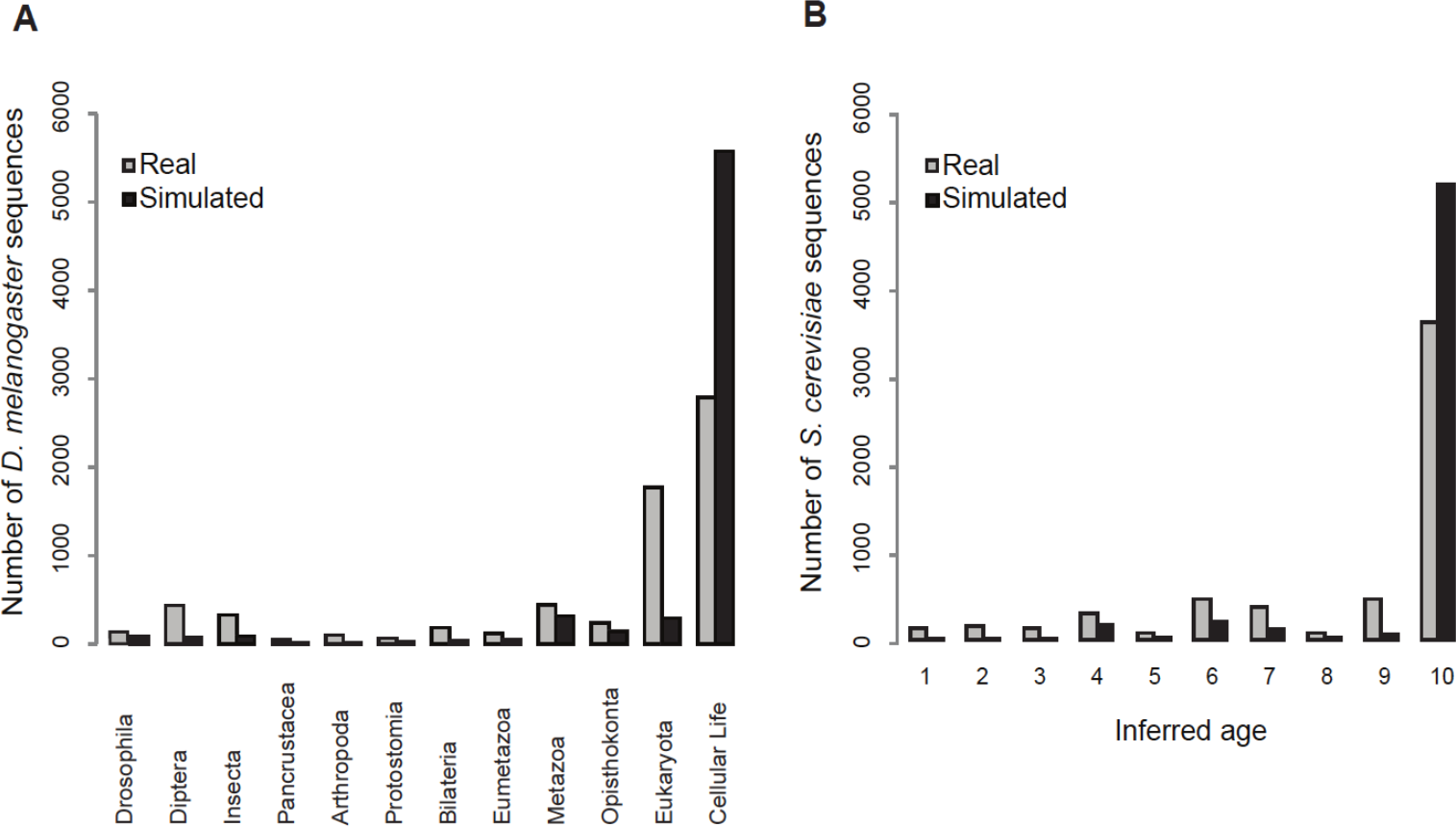
Distribution of phylostratigraphic assignments for simulated versus real sequences for *D. melanogaster* and *S. cerevisiae*. Distributions show that the majority of phylostratigraphy-based young age assignments cannot be attributed to BLAST limitations. (A) Phylostratigraphic assignments for the subset of *D. melanogaster* sequences chosen by Moyers and Zhang (2015) using real and simulated sequences; the simulated results represent the average number of sequences assigned to each phylostrata over 10 runs. (B) Distribution is redrawn from Figure 1B in Moyers and Zhang (2016)(2016), using a linear scale, rather than a log scale. Numbers indicate groups of *S. cerevisiae* ORFs of increasing conservation level within the Ascomycota, from *S. cerevisiae*-specific (1) to conserved *in S. pombe* (10).

We next investigated whether the number of simulation runs performed would influence the comparatively large numbers of young sequences found in real versus simulated data (Figure 1). Indeed, since simulations are by nature stochastic, the list of sequences found error-prone in a given simulation run is expected to vary somewhat each time a new simulation run is performed. Therefore, the number of sequences found error-prone could potentially increase towards values equal or superior to the values observed in real data if the union of multiple simulation runs was considered. To evaluate the impact of the number of simulation runs on the estimated BLAST false negative rate, we performed a saturation analysis on a series of 10 independent runs simulated by Moyers and Zhang (2015) on *Drosophila* sequences. Starting from 3,840 sequences found to lack a remote homologue at the cellular life phylostrata in the real phylostratigraphy, we asked how many of these sequences are found susceptible to BLAST limitations in the union of up to ten successive independent simulation runs. We found that, while on average a single simulation run identifies 866 error prone sequences, this number increases only to 1,006 when the union of ten simulation runs is considered (Figure 2). The number of simulation iterations thus barely affects estimates of BLAST limitations. Therefore, while phylostratigraphic methods should be improved to reduce an already low false negative rate of 5-15%, technical BLAST artifacts cannot explain away the punctuated evolution of protein coding genes nor the evidence for *de novo* emergence throughout evolutionary time.

**Figure 2:**
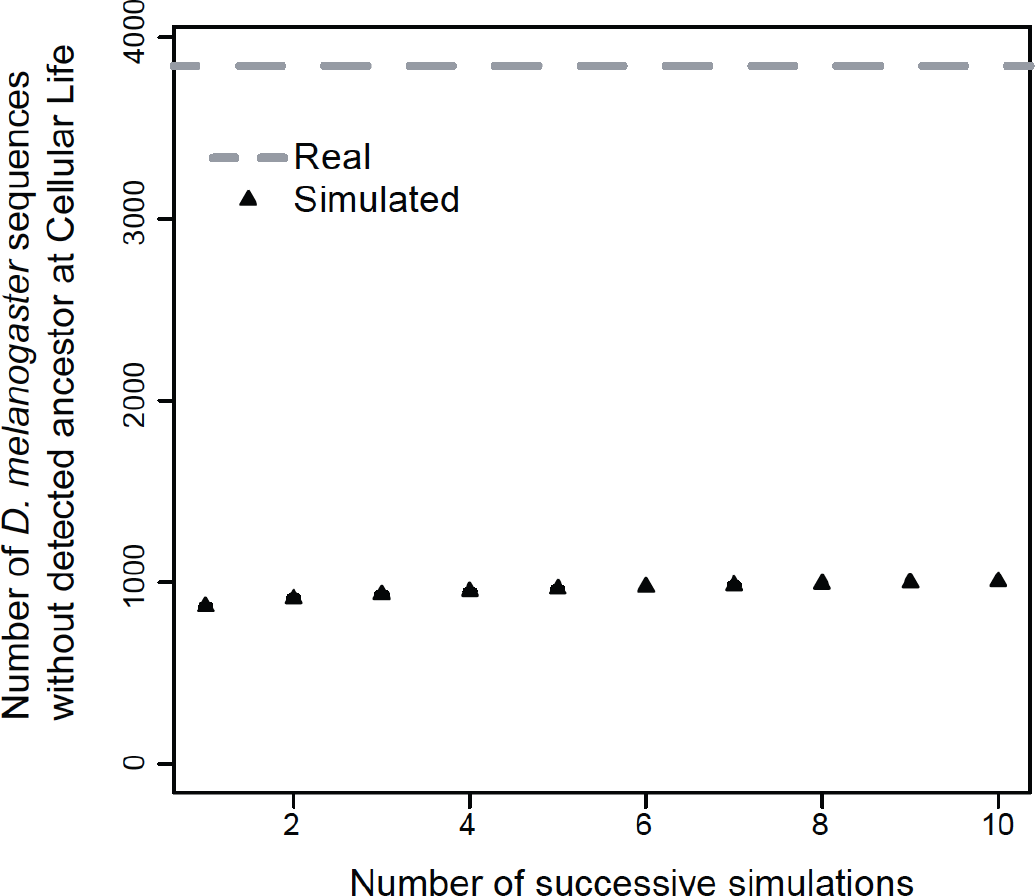
The number of *Drosophila melanogaster* genes classified young using real data (dashed grey line) that are also found susceptible to BLAST limitations by Moyers and Zhang’s simulations (2015) (black triangles) saturates rapidly. The average of 15 random permutations of 10 successive simulations is shown; standard errors of the mean are not shown because they are shorter than the height of the triangles.

### No "spurious"patterns of phylostratigraphy

We next asked if BLAST limitations, whatever their magnitude, could have influenced the published correlations between phylostratigraphic and biological patterns. This was attempted by Moyers and Zhang (2015) and Moyers and Zhang (2016) who, although they admittedly could not reproduce the exact patterns that were found in real data, claimed that the simulated sequences also yielded evolutionary patterns that appear interesting and significant, and that one would have no possibility to tell which ones are correct. Specifically, Moyers and Zhang criticize three series of results that we previously published: 1) Domazet-Lošo et al. (2007) reported that the genes expressed in ectoderm, mesoderm and endoderm during *Drosophila* development show a non-random distribution of phylostratigraphic conservation levels; 2) Domazet-Lošo and Tautz (2008) showed that human disease genes tend to be more ancient than expected; 3) Carvunis et al. (2012) and Abrusán (2013) found that many structural and functional characteristics of ORFs sequences (such as length, expression level and hydropathicity) correlate with their date of emergence in the Ascomycota fungal phylogeny. Moyers and Zhang (2015) also re-investigated the finding that new gene origination peaked in the common ancestor of Bilateria (Domazet-Lošo et al. 2007) but they could not recapitulate this pattern in their simulations.

First, we re-examined the claim according to which simulated *D. melanogaster* sequences may yield significant over-and underrepresentation of genes from certain age groups that are expressed in ectoderm, mesoderm and endoderm (Moyers and Zhang 2015). We obtained the simulated sequence sets from the authors and reproduced the patterns shown in Moyers and Zhang (2015). However, we could not reproduce the corresponding significance values, (Figure 3A), even without Bonferroni correction (supplementary table S2) indicating some problem in Moyers and Zhangδs significance calculations. Moyers and Zhang (2015) have combined 10 simulation runs to obtain this pattern and its significance, but we show that the individual runs have no common trend and that none are significant (Figure 3B). In fact, the magnitude of the log-odds ratio obtained in their simulations (Figure 3A) is much lower than reported in the original study (Domazet-Lošo et al. 2007).

**Figure 3:**
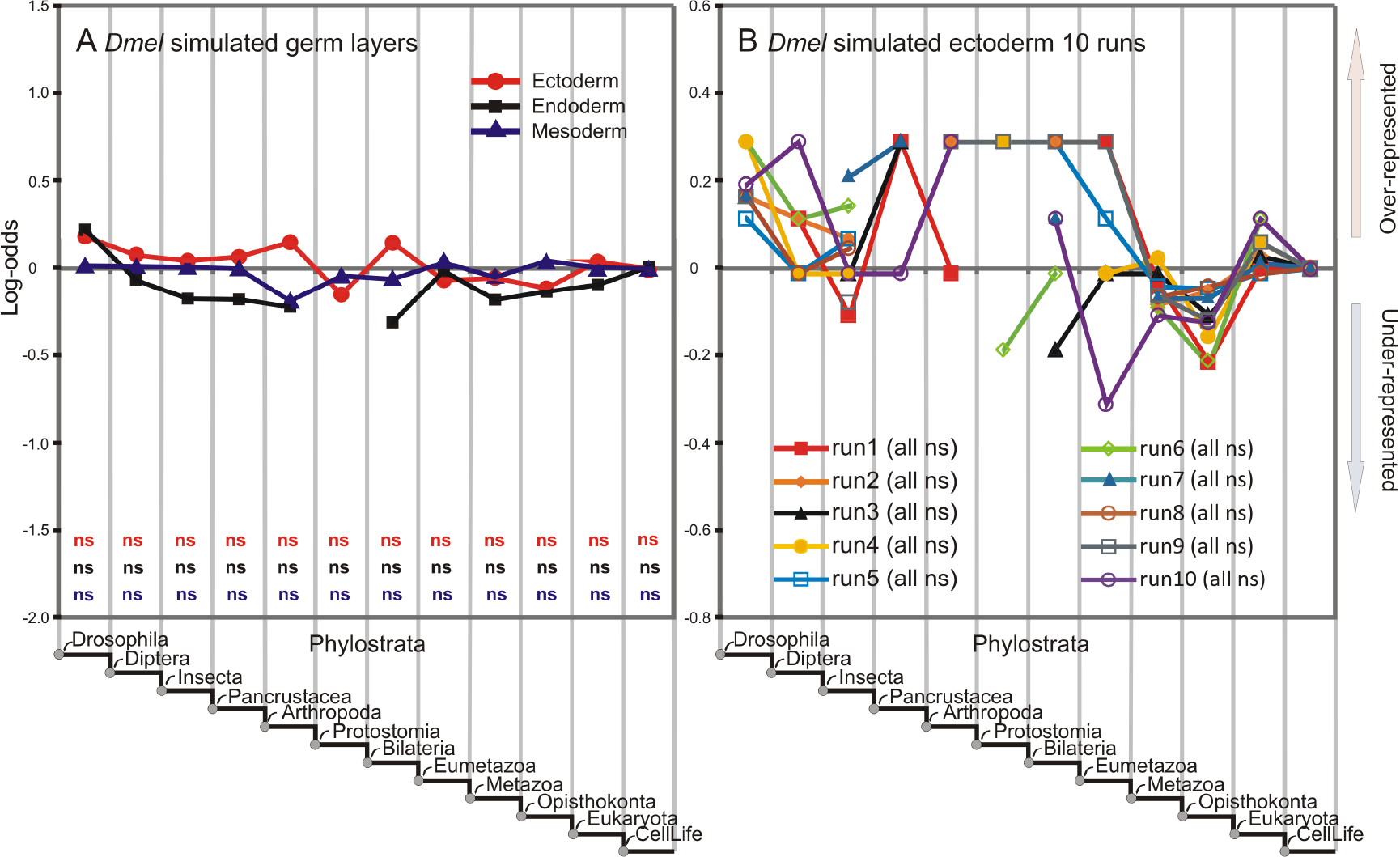
Recreated phylostratigraphic analyses of gene expressions in simulated germ layers from Moyers and Zhang (2015). (A) Overrepresentation profiles averaged over 10 simulated datasets reported by Moyers and Zhang (2015) in their figure 3c. In contrast to the claims made by Moyers and Zhang (2015), none of the deviations is significant by hypergeometric test (ns) with Bonferroni correction. (B) Overrepresentation profiles in ectoderm for 10 replicated simulations. Note the instability of profiles across the replicates and number of phylostrata without any expressed genes. None of the deviations at any phylostrata is significant by hypergeometric tests (ns).

Nevertheless, to further evaluate the robustness of the central original finding that the genes emerging after the origin of eukaryotes tend to be expressed more in ectodermal than in endodermal and mesodermal tissues (Domazet-Lošo et al. 2007) we repeated the analysis of *Drosophila* germ layers using the most recent expression and sequence databases. The input dataset we use here was much better populated compared to the datasets in the original study (see Methods). This analysis confirmed the initial finding that ectoderm is expressing more evolutionary younger genes compared to the mesoderm and endoderm (Figure 4A). However, some of the fluctuations seen in the original data (i.e. Fig. 2a in (Domazet-Lošo et al. 2007) appear to be more smoothed out in the current analysis, likely due to the more extensive data available. When we removed from the analysis genes that Moyers and Zhang found susceptible to the BLAST error in their simulations (192 out of 4157 genes with expressions) the general profiles remained largely unaffected (Figure 4B), i.e. such potentially misplaced genes do not distort the major results.

**Figure 4:**
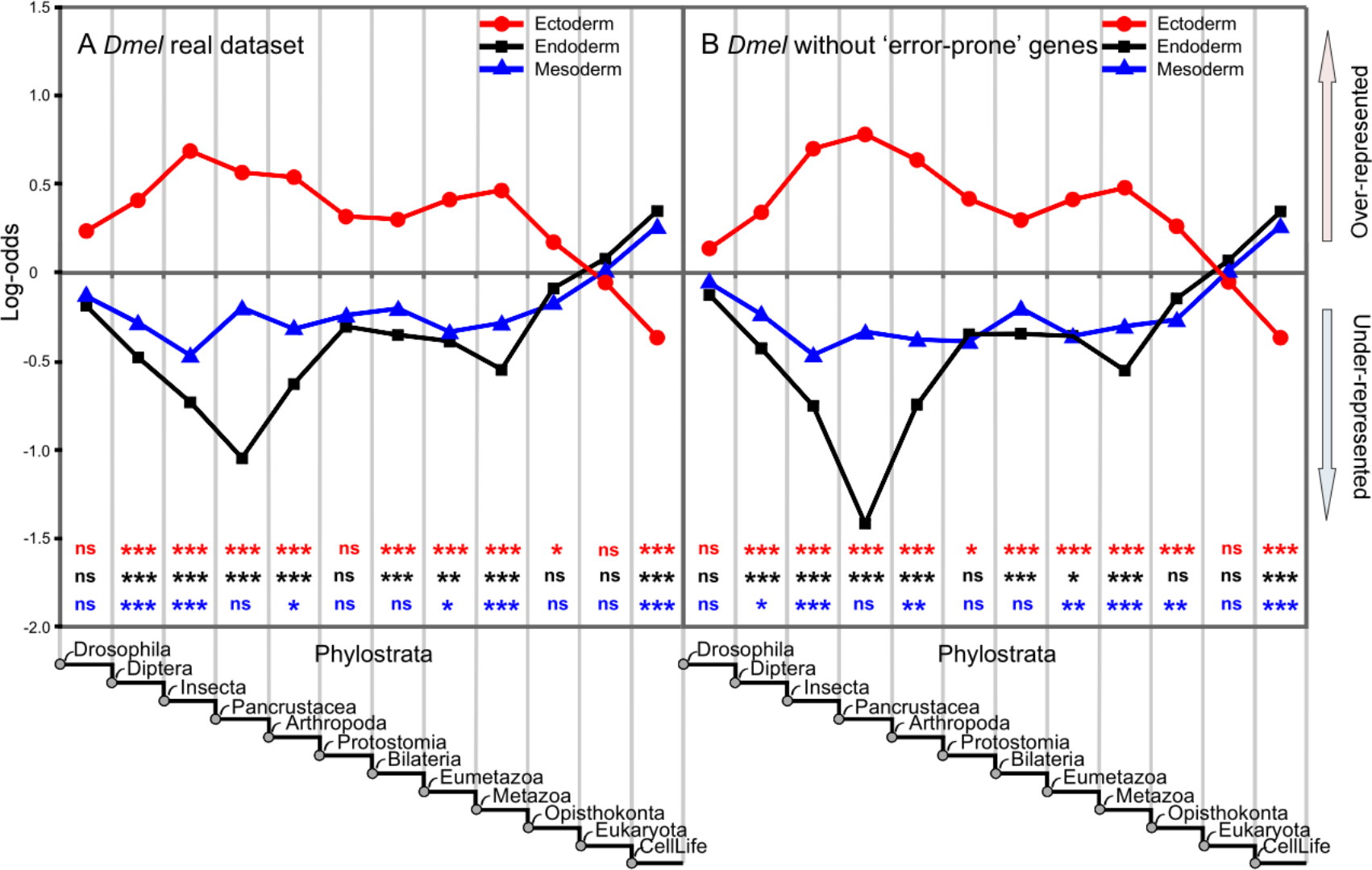
Updated phylostratigraphic analyses of gene expressions in fruit fly germ layers from Domazet-Lošo et al. 2007. (A) Real phylostratigraphic map using the latest sequence and expression databases. (B) Real phylostratigraphic map after the removal of genes that are found to be susceptible to BLAST limitations by Moyers and Zhang (2015). Note that the profiles remain largely unaffected. Stars represent significances after hypergeometric test with Bonferroni correction (^*^ at 0.05 level, ^**^ at 0.01 level and ^***^ at 0.001 level).

Second, we observed another statistical problem in Moyers and Zhang’s (2015) critique of our finding that human disease genes are enriched in ancient genes relative to young ones, which was originally shown by assessing the significance of log-odds ratio per phylostratum (Domazet-Lošo and Tautz 2008). In Moyers and Zhang’s analyses, a set of human genes was simulated and they reported "a positive correlation between the inferred age of a gene and its probability of being a disease gene (Spearman’s ρ = 0.623, P = 0.004; Fig. 4)." (quote from (Moyers and Zhang 2015). This statement is actually different from our finding that two phylostrata (origin of life and origin of metazoans) show a significant enrichment of disease genes and that young genes are significantly under-represented (Domazet-Lošo and Tautz 2008). In fact, given that Moyers and Zhang’s simulation framework assumes that all genes, including all disease genes, are old, their analysis could not have returned any other result than that the disease genes tend to be old. Indeed, the oldest phylostratum is an attractor where they placed all genes, including all disease genes. With this input constraint, it is very hard to produce conditions, by any realistic simulation, that would return the result that disease genes have no tendency to be old. Hence, Moyers and Zhang’s correlation analysis was circular.

Third, we investigated whether the evolutionary continuum of structural and functional features in fungal ORFs reported by Carvunis et al. (2012) and Abrusán (2013) could be attributed to false negatives in BLAST, as claimed by Moyers and Zhang (2016). In the original study, Carvunis et al. (2012) had included several controls to show that the observed correlation between conservation level and ORF length was not an artifact of BLAST errors. They showed that a significant correlation could be reproduced even when limiting analysis to ORF sequences with BLAST hits covering at least 80% of sequence length, and they implemented a series of partial correlations to control for the known crosscorrelations between length, expression level and evolution rates. Furthermore, all correlations reported by Carvunis et al. (2012) were checked for robustness by verifying that significance was also observed when excluding very young ORFs and when sampling only 50 ORFs from each phylostratum (100 bootstrap simulations per correlation statistics). Moyers and Zhang (2016) did not reproduce any of these controls.

To determine whether BLAST errors as estimated by Moyers and Zhang (2016) may nevertheless explain the observed evolutionary continuum, we revisited the original analyses after excluding all ORFs deemed to be susceptible to the BLAST artifact by Moyers and Zhang (2016). We calculated the Kendall correlations between various sequence features and evolutionary age, where age was inferred in three different ways: by phylostratigraphy on simulated ancestor sequences (as performed by (Moyers and Zhang 2016)), by phylostratigraphy on the real sequences (as performed by (Carvunis et al. 2012)) of the same 5,878 ORFs included in the simulations, and by phylostratigraphy on the real sequences (as performed by (Carvunis et al. 2012)) of 5,209 ORFs shown to be robust to the BLAST false negative errors by Moyers and Zhang’s simulations (i.e. after removing all ORFs found to be young in the simulations). All trends reported by Carvunis et al. (2012) were qualitatively the same and statistically robust to the BLAST false negative rate, as they held up after removing the subset of ORFs susceptible to have been erroneously classified as young in the original study (Figure 5, Table 1). Hence, rather than undermining the original conclusions, the simulation approach actually strengthens them.

**Figure 5:**
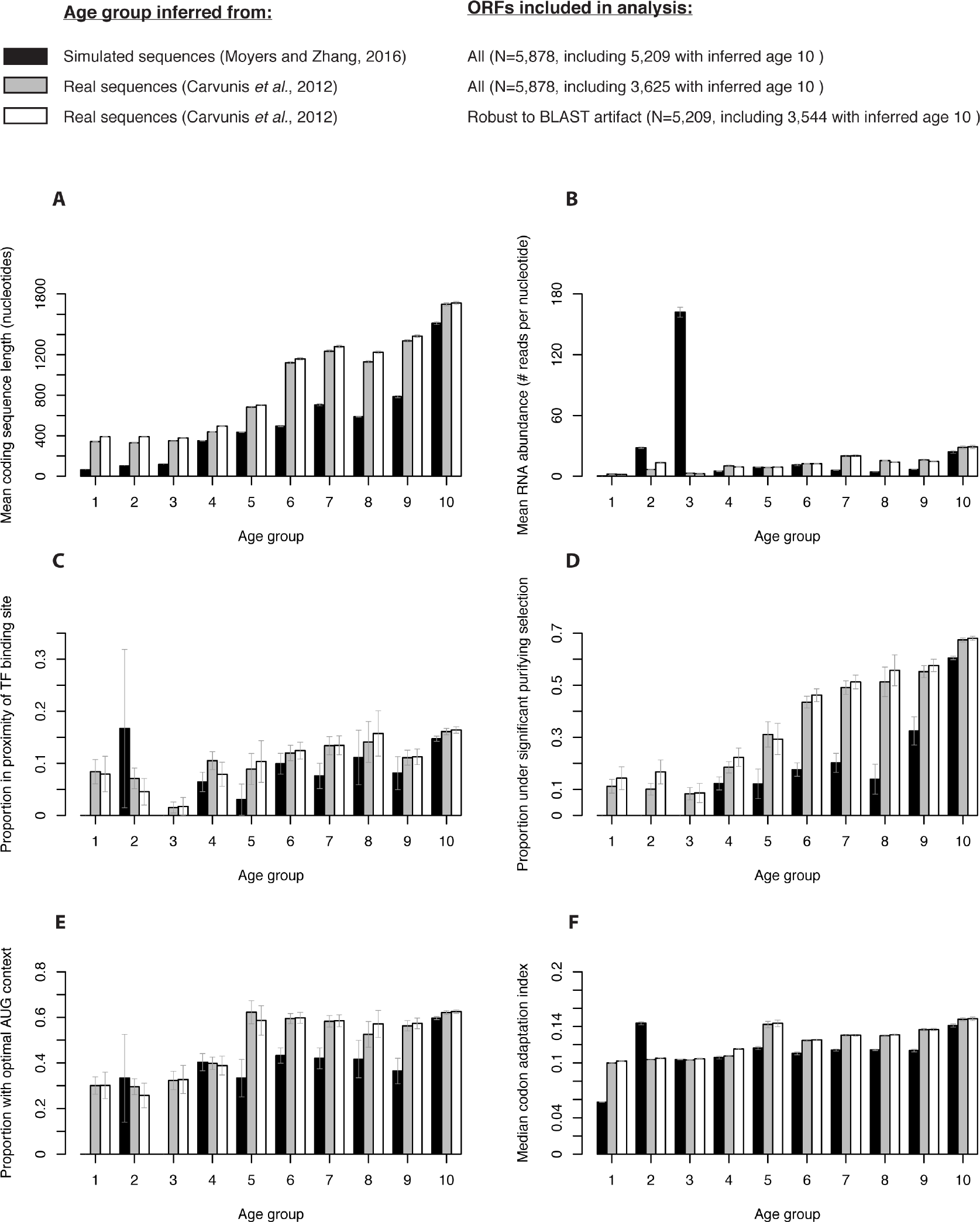
Distribution of six biological features for 5,878 *S. cerevisiae* ORF sequences with age inferred from real data (grey), for the same 5,878 ORF sequences with age inferred in simulations (black) and for 5,209 ORF sequences shown to be robust to potential BLAST artifact because they are assigned to the oldest age group in the simulation, with age inferred from real data (white). Vertical error bars represent standard error of the mean (A and B), standard error of the proportion (C, D and E) or standard error of the median (F).

**Table 1.** Correlations (Kendall’s tau) between inferred ORF age and various biological features.

| Comparison | ORF Length | RNA abundance | Proximity to TFBS | CAI | Purifying selection | Optimal AUG context |
| --- | --- | --- | --- | --- | --- | --- |
| All real proteins | 0.39** | 0.26** | 0.08* | 0.31** | 0.32** | 0.13** |
| Real proteins, robust to BLAST artifact | 0.33** | 0.17** | 0.07* | 0.25** | 0.23** | 0.09* |
| Simulated proteins | 0.28** | 0.26** | 0.06* | 0.21** | 0.27** | 0.12** |

\* $P < 0.05$ . \*\* $P < 1E-16$ . Note that the conservation levels in the original Carvunis et al. (2012) paper and the first half of Table 1 from Moyers and Zhang (2016) comprised level 0, which corresponds to non-annotated *S. cerevisiae* ORFs, plus levels 1 to 10, estimated by phylostratigraphy on real or simulated sequences. Here only levels 1 to 10 are considered.

We next sought to understand why the simulation approach yielded results that were somewhat comparable to the real data. As mentioned above, in their simulations, Moyers and Zhang (2015); Moyers and Zhang (2016) started with the real sequences - rather than *in silico* generated random sequences - and let them evolve randomly according to rate parameters inferred from real alignments among closely related species. Hence, the true features of these sequences are inherently still implied in the model, i.e. the same sequences that are short or fast evolving in reality are also short or fast-evolving in the simulations. These sequences in turn are most likely to be misclassified in the simulations since length and evolution rate affect the performance of BLAST. This leads to circularity, since it has been well established that recently emerged genes are short and evolve rapidly, in part through studies of closely related species where no BLAST error could reasonably be invoked (Reinhardt et al. 2013; Palmieri et al. 2014; Ruiz-Orera et al. 2015).

The effect becomes very evident when one looks at the overlap between the real sequences placed at particular nodes and the simulated equivalents (Figure 6). Of 6,629 *D. melanogaster* sequences and 5,878 *S. cerevisiae* sequences with ages inferred both in the original phylostratigraphies and in simulations, only 1,324 and 699 appear susceptible to BLAST error for *D. melanogaster* and *S. cerevisiae*, respectively. The vast majority of these sequences (76% and 88% for *D. melanogaster* and *S. cerevisiae*, respectively) were also assigned a young age group in the original phylostratigraphies. Given these overlaps, it is evident that the characteristics of sequences of any given age group will be somewhat comparable between simulated and real data, since the simulated data comprises mostly of sequences in the real data, with noise added and without dissociating the age-influencing features (length and divergence rate) from other features such as expression level. It is this circularity, rather than the false negative rate of BLAST *per se* (the alleged ‘phylostratigraphy bias’), that leads to seemingly similar patterns in the real and simulated data. If one wanted to assess whether the false negative rate of BLAST *per se* would give rise to such significant patterns, one should randomly distribute the rate parameters across genome sequences to simulate their evolution along the phylogeny in a manner that would be independent of their true features.

**Figure 6:**
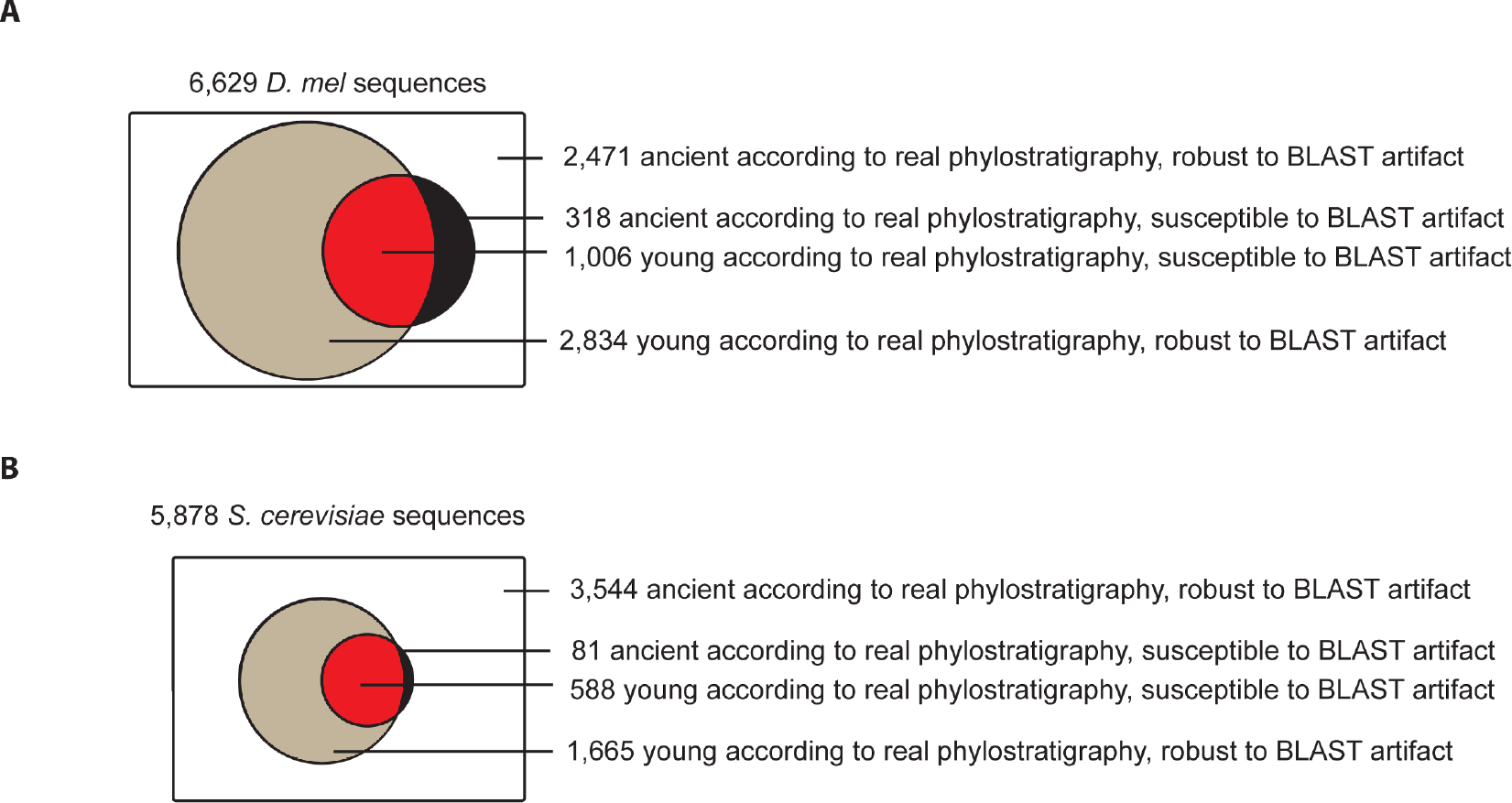
Pie charts representing sequences in the real phylostratigraphy and their relation to the BLAST artifacts in the Moyers and Zhang simulations for *D. melanogaster* (A) and *S. cerevisiae* (B). The majority (74%) of sequences found young in real data are robust to BLAST artifact (grey). Some sequences are found ancient in the real data but not in the simulated data (black), indicating that the phylostratigraphic methods used in the real data were more sensitive than those used on the simulated data. The only sequences whose phylostratum of origin may be underestimated are in red. For *Drosophila*, a conservative approach was taken where we counted as susceptible to BLAST artifact all sequences found young in at least one of 10 simulation runs. For yeast, a single run was performed and analyzed. Note that the proportion of sequences found young is larger in *Drosophila* (A) than in yeast (B) because the species tree considered is much deeper.

### Phylostratigraphy and de novo evolution

As already pointed out above, Moyers and Zhang (2016) address the question of *de novo* evolution only rather indirectly. Since it is generally acknowledged that strong evidence for *de novo* evolution can only be derived from shallow phylogenies where the non-coding sequences that gave rise to the newly evolved genes can still be traced, it is of less relevance to test BLAST error in deep phylogenies. Instead, one should focus on the youngest age classes, for example the first three in the Carvunis et al. (2012) paper. Moyers and Zhang (2016) find 14 misplaced hits in the yeast simulated data, while there are 445 in the real data, i.e. over 30 times more. An unbiased observer would consider this as strong evidence for the reported notion of a particularly high emergence of putative *de novo* genes in young phylostrata (Tautz and Domazet-Lošo 2011). But Moyers and Zhang (2016) prefer to discuss this away by claiming that they have underestimated the simulated divergence by not applying a covarion model. However, as pointed out above, a covarion model with realistic parameters would not change these numbers much.

Moyers and Zhang (2016) propose an additional criterion for credible *de novo* evolution, namely a requirement to show signs of purifying selection. However, this is not seen as a strict requirement by others in the field, for two reasons. First, some investigators have regarded recently evolved ORF sequences lacking selection signatures as interesting entities representing intermediate “proto-gene” stages (Carvunis et al. 2012) that may harbor valuable information to study the mechanisms leading to the emergence of new genes even if they are not *per se* functional. These may be part of a proposed life cycle of sequences switching between a stochastic and an adaptive phase (Neme and Tautz 2014). Second, even when the interest lies in identifying adaptively evolving canonical genes of recent evolutionary origin, classic dN/dS tests of purifying selection are not considered efficient since very young *de novo* genes typically do not have acquired enough new mutations to allow these tests to become significant.

In the case of *S. cerevisiae*, Moyers and Zhang (2016) re-analyzed 16 ORFs found to be *S. cerevisiae-specific* and to show evidence of selection in the original study by Carvunis et al. (2012). Moyers and Zhang (2016) argue that these ORFs are not species-specific and lack evidence of selection. However, the methodologies used to determine species-specificity and estimate selection are very different between the two studies. Of the 16 sequences, 15 partially overlap a more conserved gene on another reading frame (overlaps are frequent in the compact yeast genomes). Carvunis et al. (2012) concentrated on the regions of these ORFs that were free from overlap and found them to be *S. cerevisiae-*specific. In contrast, Moyers and Zhang (2016) consider the full-length ORF sequences and found them to be more conserved. This is not surprising, since the full-length sequences include sequences pertaining to other genes on alternative reading frames that are indeed more conserved.

To estimate the strength of selective pressures, Carvunis et al. (2012) used the full-length sequences based on the assumption that the codon-level evolution of the alternative reading frames would not influence estimations of dN/dS on the ORF sequences of interest. Moyers and Zhang (2016) challenged these assumptions and focused this time on the overlap-free regions of the sequences to re-estimate dN/dS. However, they report only between 0-3 SNPs per region. These low numbers prevent any statistical assessment of whether the number of non-synonymous SNPs compared to the synonymous ones is more or less than what is expected under neutrality using a Fisher test. They are aware of this limitation by stating in the discussion "Nonetheless, we recognize that statistical tests of natural selection may be powerless for species-specific genes because only intraspecific polymorphism data may be used and because newly created *de novo* genes may be short." We note that this is very different from the claim made in the abstract that "there is no evidence of purifying selection on very young *de novo* genes". The correct statement should have been that there is not enough statistical power to determine the amount of purifying selection based on the nonoverlapping regions of these recently-evolved ORFs.

Moyers and Zhang (2016) concede that ‘nothing is wrong with the theoretical model of *de novo* gene birth’. Their fundamental point of contention with Carvunis et al. (2012) and Abrusán (2013), which goes beyond a mere supposed 11% false negatives, is that the original publications did not explicitly state why the observed trends would be expected from the *de novo* gene birth model. For example, Moyers and Zhang wonder ‘why the refinement of biological function of an ORF has to occur by increasing the ORF length rather than by decreasing the length’, why ‘the mean hydropathicity should decrease’ etc. They are particularly surprised to see that many of the trends continue even for older phylostrata, ‘as if the maturation of *de novo* genes takes more than 500 Myrs’. Let us here clarify these questions.

The prediction of the proto-gene model for *de novo* gene birth is actually broader than any single descriptor of genes such as length or hydropathicity: it is that the functional and structural characteristic of ORFs should follow an evolutionary continuum between non-genic sequences and genes (Carvunis et al. 2012). For example in the case of *S. cerevisiae*, non-genic sequences are rippled with short ORFs thought to appear and disappear by chance through neutral mutations. In contrast, canonical protein coding genes with established biological functions are on average much longer. Thus, the continuum prediction of the *de novo* gene birth model is that, in Ascomycota, ORF length should increase on average with evolutionary conservation. This is not meant to imply that ORF length would continuously increase, for all ORFs, over extended periods of evolutionary time. Rather, the statement simply indicates that, since the randomly appearing ORFs are virtually all short, only those that have been maintained over longer periods of time can be long, which leads to an increase of average length over time-since-emergence. This trend is indeed also seen in studies of vertebrate taxa (Toll-Riera et al. 2009; Neme and Tautz 2014). One could imagine that the continuum prediction would actually predict the opposite trends in species where randomly appearing ORFs would tend to be longer, as may be the case in the Mycoplasmataceae lineage, which uses only two stop codons (Tatarinova et al. 2016). Because the continuum prediction is so general, it allows investigators to discover evolutionary trends without a priori suppositions of how *de novo* proteins should evolve. Rather, the data can be examined with an open mind thanks to the power of phylostratigraphy.

Moyers and Zhang (2016) discuss also whether there is a prevalence of origination of new genes via gene duplication or *de novo* evolution. Carvunis et al. (2012) have discussed long-term trends and concluded that *de novo* evolution may be more frequent. This was also the finding of Neme and Tautz (2014) in vertebrates. The overall pattern of extensions of transcript length, number of exons, length of ORFs and acquisition of domains makes it more likely that new genes are initially short. If one would want to explain such trends through a duplication-divergence model, one would have to assume either that short genes are more likely to be duplicated, or that genes become shorter after duplication. Neither of these trends have so far been reported.

## Conclusion

The studies by Moyers and Zhang have revisited previously discussed important issues, but have failed to provide much new insights or evidence for the existence of a hypothetical phylostratigraphic bias due to the use of BLAST. Still, for genes that have arisen at deep phylogenetic nodes, there will be some uncertainty whether they evolved according to the duplication-divergence or the *de novo* evolution model. But as pointed out in the introduction, this is of secondary importance for tracing evolutionary patterns through phylostratigraphy. For genes that have arisen very recently, there is now overwhelming evidence that *de novo* gene birth has occurred repeatedly in many lineages, where possible deficiencies of detection via BLAST play no practical role. There is no reason to assume that the proven high rate of *de novo* evolution of transcripts has not occurred throughout evolutionary history. Although the turnover of *de novo* genes seems very high (Palmieri et al. 2014; Neme and Tautz 2016), some will inevitably have been retained, in particular at times of major radiations and evolution of new lineages (Tautz and Domazet-Lošo 2011). We concur with Moyers and Zhang´s (2016) suggestions that gene by gene studies will provide deeper insights into these questions and that phylostratigraphic methodologies could be further improved to date the emergence of sequences with even higher accuracy. Research in this direction should consider not only BLAST false negatives, where sequences appear young versus ancient and fast evolving, but also false positives, where BLAST hits are spurious versus true homologues of the query sequences. However, we refute the conclusion that genome-wide evolutionary trends are "insufficient and confounded by phylostratigraphic error." Instead, the data presented here demonstrate unequivocally that phylostratigraphic analyses of patterns of gene emergence and evolution are robust to the false negative rate of BLAST, whether it is in the range of 5% or 15%. Finally, the errors we detected in the Moyers and Zhang´s (2015) analyses urge for the careful use of statistics when testing phylostratigraphic patterns.

## Methods

### Reanalysis of Moyers and Zhang 2015 dataset and statistics

Moyers and Zhang kindly sent us their dataset with list of genes that contained ectoderm, endoderm and mesoderm and their simulated phylostrata over 10 simulation runs. For our saturation analysis (Figure 2), we generated 15 random permutations of these 10 simulation runs. For each permutation, we calculated the number of *Drosophila melanogaster* genes found young in the real phylostratigraphy (lacking a detected ancestor at Cellular Life) that could have been misplaced when considering the union of 1 simulation, 2 simulations, …, 10 simulations. We then averaged the numbers over the 15 random permutations. We also repeated their ontogeny analysis and calculated hypergeometric tests with Bonferroni correction for all 10 runs and three germ layers (supplementary table S2). We created our Fig. 3A to match their Fig. 3C by using average values of 10 runs. To be able to calculate significances by hypergeometric tests we rounded rational numbers obtained by averaging to integers.

### Phylostratigraphic reanalysis of the expressions in the fruit fly germ layers

To allow broad sequence similarity searches we first built a custom built protein database by combining complete genomes from National Center for Biotechnology Information (NCBI), Ensembl and Joint Genome Institute (JGI). In total we collected 113,834,351 protein sequences from 25,223 genomes. To reduce large redundancy of prokaryotic sequences (23,675 prokaryotic genomes) we clustered prokaryotic parts of the database with the CD-HIT at 90% identity (Li and Godzik 2006). After this procedure our database contained 43,899,817 protein sequences. For comparison, in the original study we used a database that comprised 2,777,855 protein sequences (only around 2% of the present database size).

We compared 13,389 protein sequences of *Drosophila melanogaster* retrieved from the Ensembl database (Yates et al. 2016) against the protein database by using the similarity search algorithm BLASTP (Altschul et al. 1997) at E-value cut-off of 1e-03 (Domazet-Lošo et al. 2007). Using the obtained BLAST output we mapped the fruit fly genes onto a consensus phylogeny (12 phylostrata) using the most-distant BLAST match above the significance threshold (BLAST E-value less than 1e-03) as described in the original study (Domazet-Lošo et al. 2007). This updated *Drosophila* phylostratigraphy is provided in supplementary table S1.

### Drosophila expression data and statistics

We retrieved *in situ* hybridization expression data for 4,157 fruit fly genes that show tissue-specific expression during ontogeny from the Berkeley *Drosophila* Genome Project (Tomancak et al. 2002). In total, this set of genes contributes to 38,627 expression domains expressed over multiple tissues and the different stages of the ontogeny. In the original study we had used 1,967 genes with 10,432 expression annotations (only around 27% of the present expression dataset). We divided the fruit fly expression dataset into subsets corresponding to the specific germ layer (either ectoderm, endoderm or mesoderm). For every germ layer we performed an over-representation analysis by comparing a frequency of expression domains in a phylostratum to a frequency in the total dataset (expected frequency) (Domazet-Lošo et al. 2007; Domazet-Lošo and Tautz 2008; Domazet-Lošo and Tautz 2010b; Šestak et al. 2013; Šestak and Domazet-Lošo 2015). Obtained deviations, i.e., more or less expression than expected, are depicted in the figures by log-odds ratios and their significance was tested by two-tailed hypergeometric tests (Rivals et al. 2007) controlled for multiple comparisons via a Bonferroni correction.

### Fungal data and statistics

The conservation levels of *S. cerevisiae* ORFs was estimated by Carvunis *et al.* (2012) and simulated by Moyers and Zhang (2016). Moyers and Zhang kindly provided us with the results of their simulations. Only 5,878 ORFs that were assigned a conservation level by both studies are included here. ORF characteristics (length, expression level etc.) were taken as in Carvunis *et al.* (2012). Distributions, error bars and p-values were computed using R scripts.

## Acknowledgements

We thank B. Moyers for discussion and for providing analysis files, as well as G. Abrusán, M. Calderwood, H. Carter, J. Castresana, B. Charloteaux and J. Kreisberg for discussion, comments and suggestions on the manuscript. We thank the following funding organizations for support of our work: TD-L: City of Zagreb and Adris Foundation grants; AC: National Institute of General Medical Sciences K99 GM108865; MA: grant BFU2015- 65235-P from MINECO/FEDER, EU; DT: ERC grant NewGenes, 322564.

